# Exploring diverse routes to high-affinity-antibody variable domains through deep-sequencing-informed machine learning

**DOI:** 10.64898/2026.05.28.728451

**Authors:** Sakiya Kawada, Tomoyuki Ito, Hikaru Nakazawa, Yoichi Kurumida, Yutaka Saito, Mitsuo Umetsu

**Author notes:** Both authors contributed equally to this work (Sakiya Kawada, Tomoyuki Ito).

## Abstract

The integration of in vitro selection, deep sequencing, and machine learning (ML) has recently been developed as a powerful strategy for discovering functional antibodies. However, how training data composition and ML search space design influence the identification of high-affinity variants remains unclear. Here, we aimed to optimize ML-integrated directed evolution for functional antibody discovery by selecting training data from deep sequencing analysis. By performing phage display selection using camelid heavy-chain antibodies (VHHs), we demonstrated that early-round data, retaining more binding-negative variants, can be superior for training models to identify high-performance VHHs. We also investigated a lead-independent ML search space design by focusing on conserved residues in final rounds, successfully identifying variants with higher affinities than those from lead-based maturation (*K*_D_ = 7.9 nM). These findings demonstrate that training data selection and search space design are critical for successful ML-guided antibody engineering and provide diverse pathways for discovering high-affinity VHH variants.

## Introduction

Antibodies are molecular recognition proteins widely used as therapeutic and diagnostic agents^1,2^. Target-specific antibodies or antibody variable domains are typically developed by animal immunization or directed evolution using in vitro selection with surface-display systems. In these surface-display systems, genetic libraries are basically constructed through random mutagenesis of antibody variable domains, and encoded variants are displayed on platforms such as phage^3^ or cDNA^4^. Each in vitro selection round generally involves the removal of non-binders, followed by the elution and amplification of target-binding variants. Typically, the frequency of target-binding variants increases with iterative rounds; however, repeated cycles can lead to the enrichment of variants with low or no binding affinity but with a high propensity for amplification. This amplification bias results in a skewed population distribution, potentially leading to the loss of high-affinity variants^5–7^.

In recent years, deep sequencing analysis has enabled the evaluation of the frequency of each variant in a library, leading to the identification of predominant variants after selection^8^. These frequency data also enable us to calculate enrichment rates, by comparing pre- and post-selection libraries^6,9,10^, or specificity rates, by evaluating selections in the presence or absence of the target antigen^7,11^. By further analyzing amino acid frequencies in selected libraries, we can identify residue positions that are efficient for affinity maturation^6,9,12,13^. In addition, frequency data can be used as performance scores in machine learning (ML) models to predict variants with improved function, thus enabling the design of an ML-guided library enriched with variants that are likely to possess desirable functions^6,7,9,12,13^. However, previous studies have reported that even the inclusion of a single variant within a given training dataset can change the ML outcomes^14^, indicating that careful evaluation of the training data preparation process may improve both the predictive accuracy of ML models and the likelihood of obtaining functional variants.

In addition to training data composition, the design of the sequence space that ML predicts (i.e., the search space) is another critical factor that determines the success of variant discovery, because ML models can rank only those variants within the search space for prediction. In ML-assisted antibody optimization, the affinity or biophysical properties are typically improved by diversifying several residues of a lead antibody, which is a target-binding variant experimentally found through in vitro selection^7,12^. For example, one previous study focused on a search space consisting of three-residue mutations around a lead antibody to optimize the binding properties^7^. However, design of the lead-based search space requires the prior experimental identification of at least one lead molecule. Moreover, if the selected lead molecule represents only a local optimum in the fitness landscape, introducing a limited number of mutations around the lead may be insufficient to discover higher-performance variants.

Here, we aimed to optimize ML-integrated directed evolution for the discovery of functional camelid heavy-chain antibodies (VHHs; Variable domains of the Heavy chains of Heavy-chain antibodies) by addressing two key factors: training data composition and ML search space design. We first developed a scaffold VHH by using consensus design based on *Camelus dromedarius* sequence data retrieved from the Single Domain Antibody Database (sdAb-DB)^15^ to ensure high structural stability. By using phage display libraries with randomized complementarity-determining region (CDR) 3 regions, we then performed in vitro selection to identify a target-binding lead VHH. To evaluate the effect of training data composition, ML models were trained by using deep sequencing data from different selection rounds and applied to lead-based affinity maturation. Our results demonstrate that early-round data, which retain a substantial proportion of binding-negative variants, can be more effective for identifying high-performance antibodies than later, more converged datasets. Furthermore, we investigated a lead-independent ML search space design by focusing on residues conserved in the final selection rounds. This approach enabled us to identify VHH variants with higher affinities than those obtained through lead-based maturation, achieving affinity in the nanomolar range (equilibrium dissociation constant, *K*_D_ = 7.9 nM). These findings demonstrate that both training data selection and search space design are critical determinants of successful ML-guided antibody engineering.

## Results

### In vitro selection using a VHH library based on a consensus scaffold

Although VHHs are inherently monomeric and structurally stable, mutations in the CDRs can compromise their stability^7^. Therefore, we developed a stable scaffold VHH by using consensus sequence design, defined as selecting the most frequent amino acid at each position from a large dataset of homologous sequences^16,17^.

The VHH derived from the consensus design (i.e., the consensus scaffold VHH) was developed on the basis of the sequence information of 260 *Camelus dromedarius* VHHs retrieved from the sdAb-DB^15^ (see Methods, Figure 1A, Figure S1A, Figure S1B, and Figure S1C). To construct the phage library, the CDR3 region of this consensus scaffold was randomized (Figure 1B). We generated four libraries with different CDR3 lengths (8, 12, 16, and 19 residues); the C-terminal residue of CDR3 was fixed as tyrosine. By using these libraries, four rounds of in vitro selection were performed against the human epidermal growth factor receptor 3 (HER3) dimer-arm peptide (Figure 1C, see Methods). The HER3 dimer-arm peptide is a region crucial for the dimerization and signaling of HER3, which is an important target in cancer therapy^18,19^. Each round consisted of (i) negative selection to remove non-specific phages (NS-eluted phages); (ii) positive selection to isolate target-bound phages (PS-eluted phages); and (iii) amplification to enrich the selected population (amplified phages). From the phage pools obtained after the third and fourth rounds of selection, 88 clones were isolated in each round, leading to the identification of two promising variants, N-14 and N-34. The *K*_D_ values were determined to be 588 nM for N-14 (CDR3: LSYVRHPY) and 2.41 µM for N-34 (CDR3: SSHTTLGAQASTRNFLTRY; Figure 1D).

**Figure 1.**
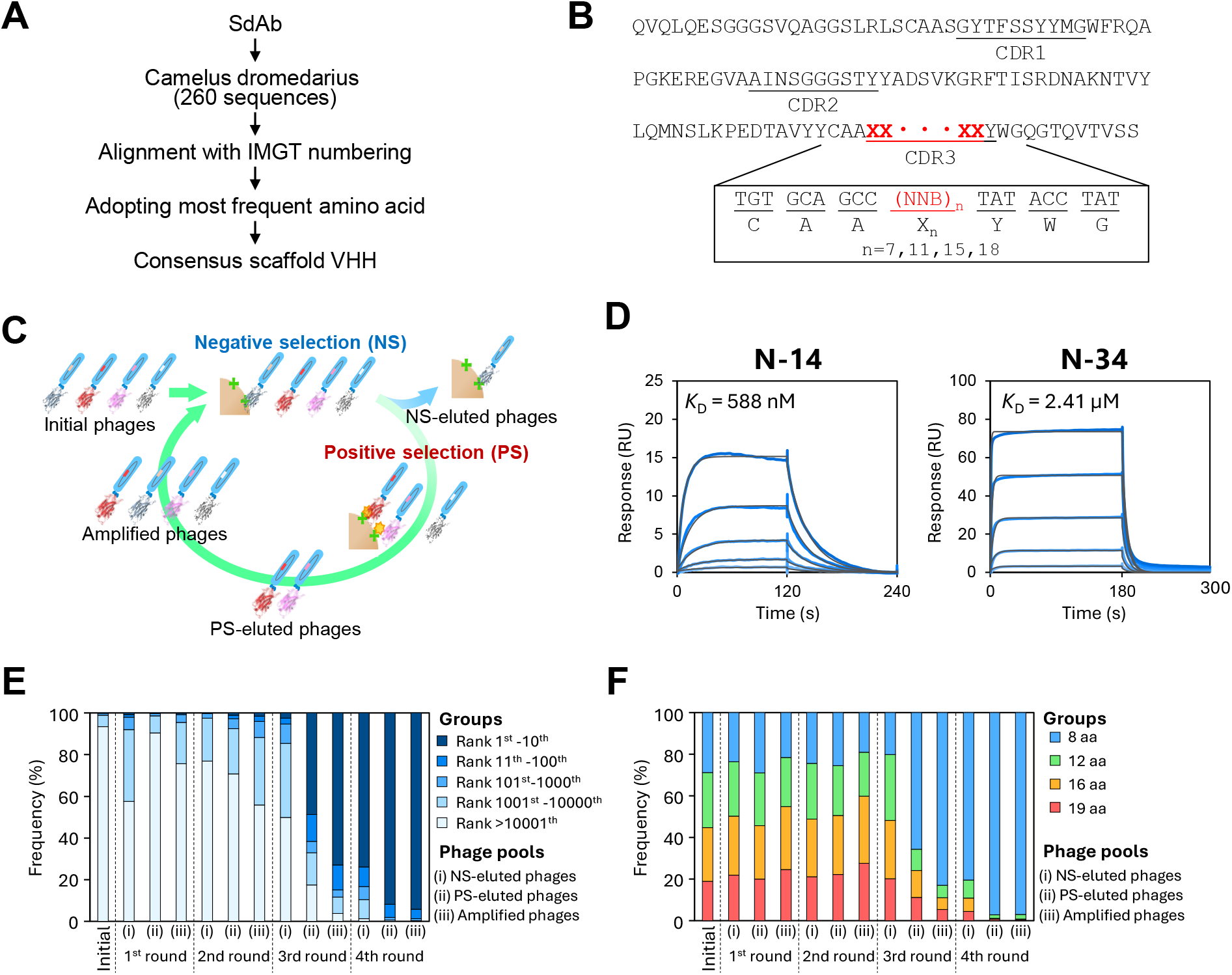
In vitro selection using a consensus-designed VHH library. (A) Workflow for consensus scaffold design. (B) Amino acid sequence of the consensus scaffold VHH. The CDR3 region was randomized with variations of 8, 12, 16, and 19 residues. The C-terminal residue of CDR3 was fixed as tyrosine. (C) Scheme of in vitro selection. Phage populations eliminated after negative selection (NS-eluted phages), those eluted after positive selection (PS-eluted phages), and the resulting amplified pools (amplified phages) were subsequently subjected to deep sequencing analysis. (D) Kinetic analysis of N-14 and N-34 by surface plasmon resonance. Three-fold dilution series were used for the analytes, starting from 1200 nM for N-14 and 20 μM for N-34 (blue lines). Equilibrium dissociation constants (*K*_D_) were determined by global fitting using a 1:1 binding model (black lines). (E) Distribution of unique sequences in (i) eluted phages from negative selection, (ii) eluted phages from positive selection, and (iii) amplified phages after selection in each round. Unique sequences within each pool were categorized by color by their frequency ranks. (F) Distribution of CDR3 lengths in each phage pool. Unique sequences were grouped by color according to their CDR3 lengths.

To further analyze biopanning, deep sequencing was performed on the initial, NS-eluted, PS-eluted, and amplified phages from each round (Figure 1C; see Methods). By calculating the frequency of each variant and tracking their sequence distribution, we observed a substantial enrichment in PS-eluted phages of the third round (Figure 1E). Notably, the proportion of variants with a CDR3 length of eight residues increased during this same step (Figure 1F). On the basis of these results, we selected N-14, a variant with an eight-residue CDR3, as the lead molecule for subsequent affinity maturation.

### Affinity maturation via ML using deep sequencing data

We previously developed a methodology for designing amino acid sequences with enhanced binding affinity by training ML models on deep sequencing data^7^. We used this approach to improve the binding affinity of N-14 (Figure S2). The performance score of each variant was defined as the ratio of the frequency in the PS-eluted phages to that in the NS-eluted phages by using data from the third round of selection, in which the strong enrichment was observed. We trained an ML model by using this sequence-score-linked data to predict the scores of variants in which four of the seven residues in CDR3 (excluding the C-terminal tyrosine) were mutated. Hereafter, the ML based on the third-round data is referred to as ML_R3. Analysis of the top 100 predicted sequences revealed that ML-proposed mutations in N-14 were concentrated predominantly at four positions (2, 4, 5, and 6), with mutation frequencies exceeding 60% at these sites (Figure 2A, Table S1). Here, the mutation frequency at a given position was calculated as the percentage of sequences that differed from the lead sequence at that position in the ML-predicted top 100 sequences. Specifically, the following amino acids showed a frequency of ≥10%: S/N/E at position 2; D/N/Q/S/E at position 4; S/H/C at position 5; and P/L/K/H at position 6 (Figure 2B, Table S2).

**Figure 2.**
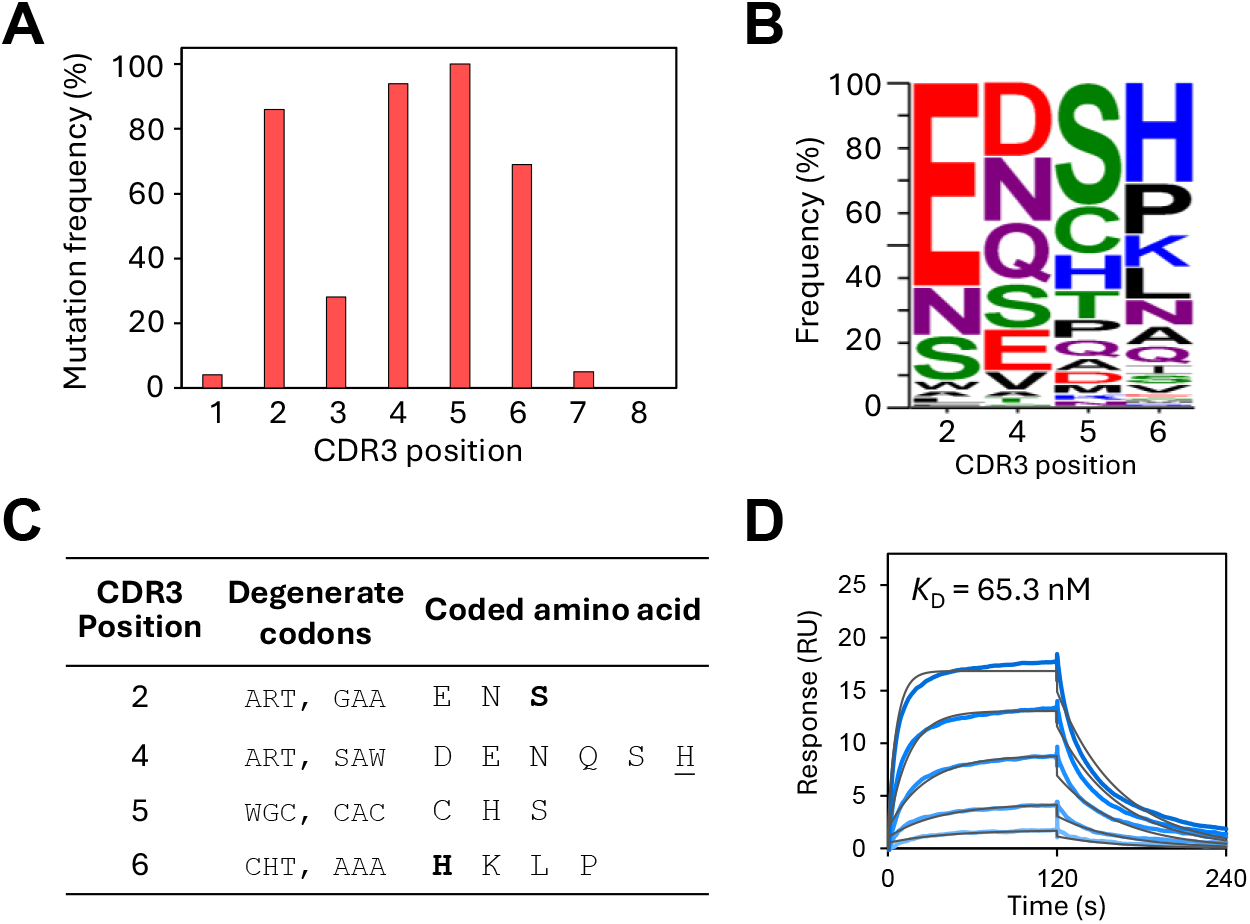
Machine learning using deep sequencing data from the third round. (A) Predicted mutation frequency at each CDR3 position of the top 100 sequences. (B) Amino acid frequencies at positions 2, 4, 5, and 6 in the ML-predicted top 100 sequences, visualized by using WebLogo. (C) Design of an ML-guided library based on amino acids that appear with frequencies exceeding 10% in the predicted top 100 sequences. The same amino acid as the one in N-14 at each position is in bold. Amino acids not present in these top 100 ML-predicted sequences are underlined. (D) Kinetic analysis of the variant 3-9A by surface plasmon resonance. Three-fold dilution series were used for the analytes, starting from 400 nM (blue lines). The *K*_D_ value was determined by global fitting using a 1:1 binding model (black lines).

On the basis of these high-frequency residues, we designed a four-residue mutated library by using degenerate codons encoding amino acids occurring at frequencies greater than 10% at each position (Methods, Figure 2C). A gene library was constructed by using the designed degenerate codons. This library was transformed into *Escherichia coli* (*E. coli*), and the binding activities of 88 randomly selected clones were evaluated. Thirty of the 88 clones (34%) had binding activities lower than one-tenth that of N-14, and 16 clones (18%) had binding activities higher than that of N-14 (Figure S3A) and contained 10 unique variants (Table S3). This indicated that, although the ML model could suggest promising candidates, it also included sequences that lacked binding activity. Among the variants with improved activity, six clones ranked within the top 10,000 by the ML model were purified by size-exclusion chromatography (SEC), and their binding affinities were evaluated using surface plasmon resonance (SPR). Three variants containing cysteine in the CDR3 had low expression levels and no detectable target binding (Figures S4A and S4B). In contrast, the remaining three variants demonstrated specific binding to the target. Notably, the most potent variant, 3-9A (CDR3: LSYHSKPY), had a *K*_D_-value of 65.3 nM (Figure 2D), representing a 9.0-fold improvement in affinity compared with that of the lead molecule N-14.

### ML using deep sequencing data from the second and fourth rounds

We investigated whether the success rate of identifying functional antibodies could be improved by using data from different selection rounds. ML models were trained using deep sequencing data from either the second or fourth round to predict the scores of four-residue variants of N-14, as well as ML_R3. Hereafter, ML based on the second- and fourth-round data are referred to as ML_R2 and ML_R4, respectively (Figure S2). To validate the models, we predicted the scores for three high-affinity variants identified in ML_R3, as well as for five variants that showed no target binding in enzyme-linked immunosorbent assay (ELISA). In the top 100 sequences predicted by ML_R2 and ML_R4, mutations were predominantly suggested at positions 4, 5, and 6 of CDR3 (Figure 3A). ML_R2 correctly distinguished three high-affinity variants above N-14, and five variants with no binding activity were ranked lower than N-14 (Figure 3B). In contrast, ML_R4 gave a poor relationship between the predicted ranks and actual binding affinities. These results suggest that the deep sequencing data from the second round may have been superior to those from the third or fourth rounds as training data for accurate prediction. The following amino acids showed a frequency of ≥10%: E/H/N at position 4; A/F/M/Q at position 5; and C/H/P/S at position 6; (Figure 3C, Table S2). On the basis of these high-frequency residues, we designed a three-residue mutated library (Figure 3D). Cysteine was excluded from the design, because CDR3 variants containing cysteine had shown low expression yields and loss of target binding (Figure S4B). We transformed this library into *E. coli* and randomly evaluated 43 clones using ELISA. We identified three variants with higher binding signals than N-14 (Figure S3B, Table S4). Notably, only four clones (9%) had binding activity less than one-tenth that of N-14. Among the three improved variants, 2-4H (CDR3: LSYHAPPY) had a *K*_D_ value of 32.8 nM (Figure 3E), representing an 18-fold higher affinity than N-14 and a 2.0-fold higher affinity than 3-9A. These results demonstrated that using second-round data not only enabled the discovery of high-affinity variants but also effectively minimized the frequency of non-functional sequences within the ML-predicted library.

**Figure 3.**
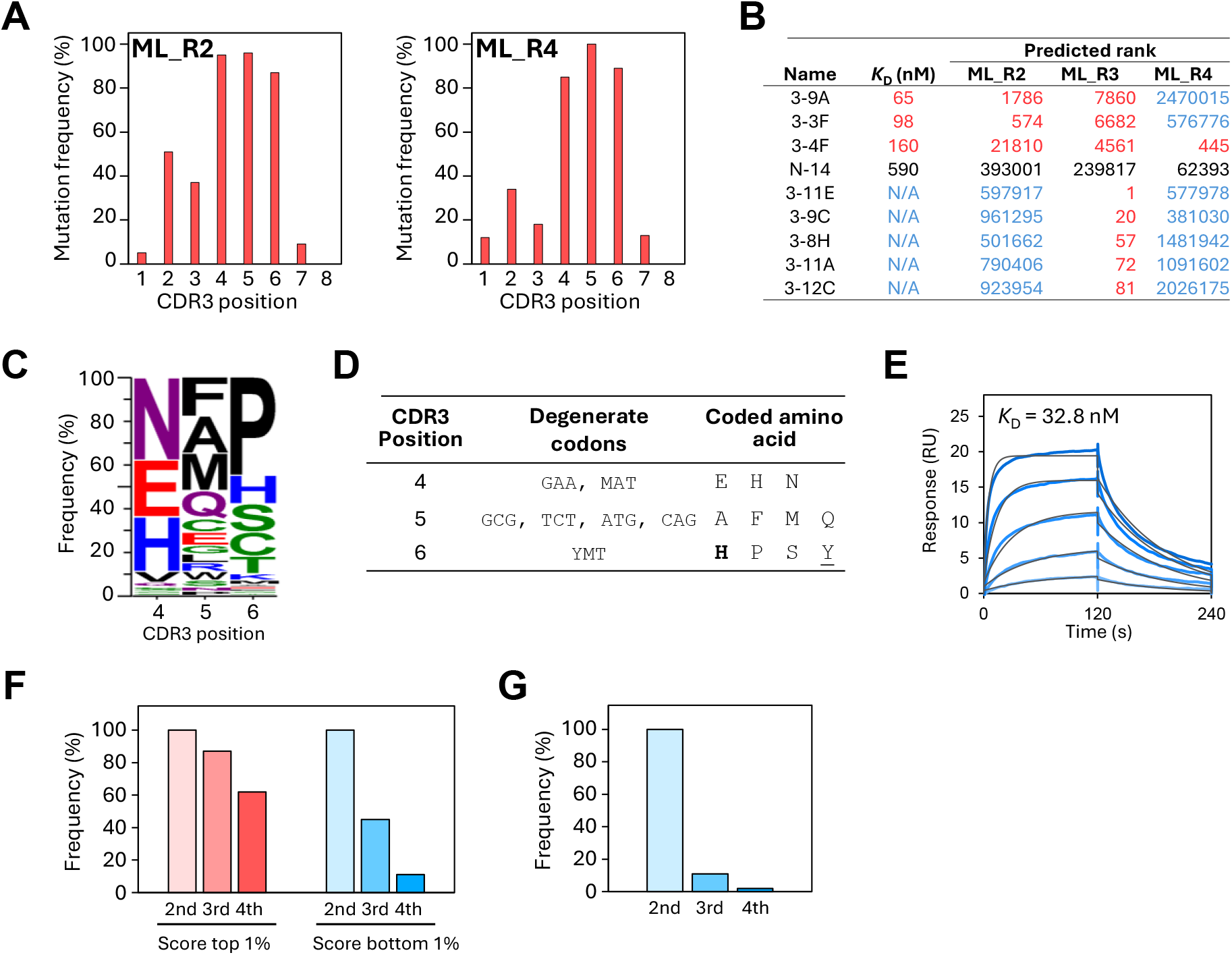
Machine learning using deep sequencing data from the second and fourth rounds. (A) Predicted mutation frequency at each CDR3 position of the top 100 sequences for models ML_R2 and ML_R4. (B) ML predicted ranks of N-14, three high-affinity variants, and five non-binding variants. Predicted ranks are compared across models (ML_R2, ML_R3, and ML_R4). Ranks superior to N-14 are in red font, whereas those inferior to N-14 are in blue. (C) Amino acid frequencies at positions 4, 5, and 6 in the ML-predicted top 100 sequences, visualized by using WebLogo. (D) Design of an ML-guided library based on amino acids that appear with frequencies exceeding 10% in the top 100 sequences predicted by ML. The same amino acid as the one in N-14 at each position is in bold. Amino acids not present in these top 100 ML-predicted sequences are underlined. (E) Kinetic analysis of variant 2-4H by surface plasmon resonance. Blue lines represent a three-fold dilution series starting from 400 nM. Black lines indicate the global fit to a 1:1 binding model. (F) Persistence of the top and bottom 1% sequences from the ML_R2 training data in the ML_R3 and ML_R4 training data. (G) Persistence of binding-negative variants identified in the ML_R2 training data within the ML_R3 and ML_R4 training data.

### Analysis of variant distribution across selection rounds

To investigate the factors that characterized the second-round data, we analyzed the behavior of the top 1% of sequences (68 variants) ranked by performance scores in the ML_R2 training data. Among these, 59 (87%) sequences were retained in the training data of ML_R3 and 42 (62%) sequences were retained in the training data of ML_R4 (Figure 3F). In contrast, we examined the bottom 1% (71 sequences) of ML_R2 training data. As a result, only 33 (45%) sequences remained in the ML_R3 training data and 8 (11%) sequences remained in the ML_R4 training data (Figure 3F). Furthermore, we focused on a subset of 1016 variants in the ML_R2 training data that were detected exclusively in the NS-eluted phage and were absent from the PS-eluted phage. Analysis revealed that only 109 (11%) of these sequences were present in the ML_R3 training data and 19 (2%) were present in the ML_R4 training data (Figure 3G). These results suggested that training datasets derived from earlier selection rounds could be preferable for antibody optimization because they provided a more balanced and informative functional landscape that included negative binders.

### Lead-independent design of sequence spaces for prediction

We focused here on the affinity maturation of N-14, a lead molecule identified through the conventional selection process. Although designing an ML search space on the basis of a specific lead molecule is effective once a lead is obtained, this approach inherently requires the prior identification of binding-positive variants. Furthermore, there is a risk that a single selected lead will represent only a local optimum within the fitness landscape. To address these limitations, we investigated the feasibility of predicting functional antibodies without a lead molecule by using deep sequencing data for both the training dataset and the design of the ML search space.

According to the deep sequencing analysis, variants with a CDR3 length of eight residues were predominant in the in vitro selection (Figure 1F); consequently, the ML search space was restricted to this eight-residue length. To further refine this space, we analyzed the amino acid frequencies of the sequences with scores of 1 or higher in PS-eluted phages in the fourth round, hypothesizing that amino acids conserved among variants in the final selection rounds contributed to binding. The frequencies of leucine at position 1 and alanine at position 7 of CDR3 exceeded 50% (Figure 4A).

**Figure 4.**
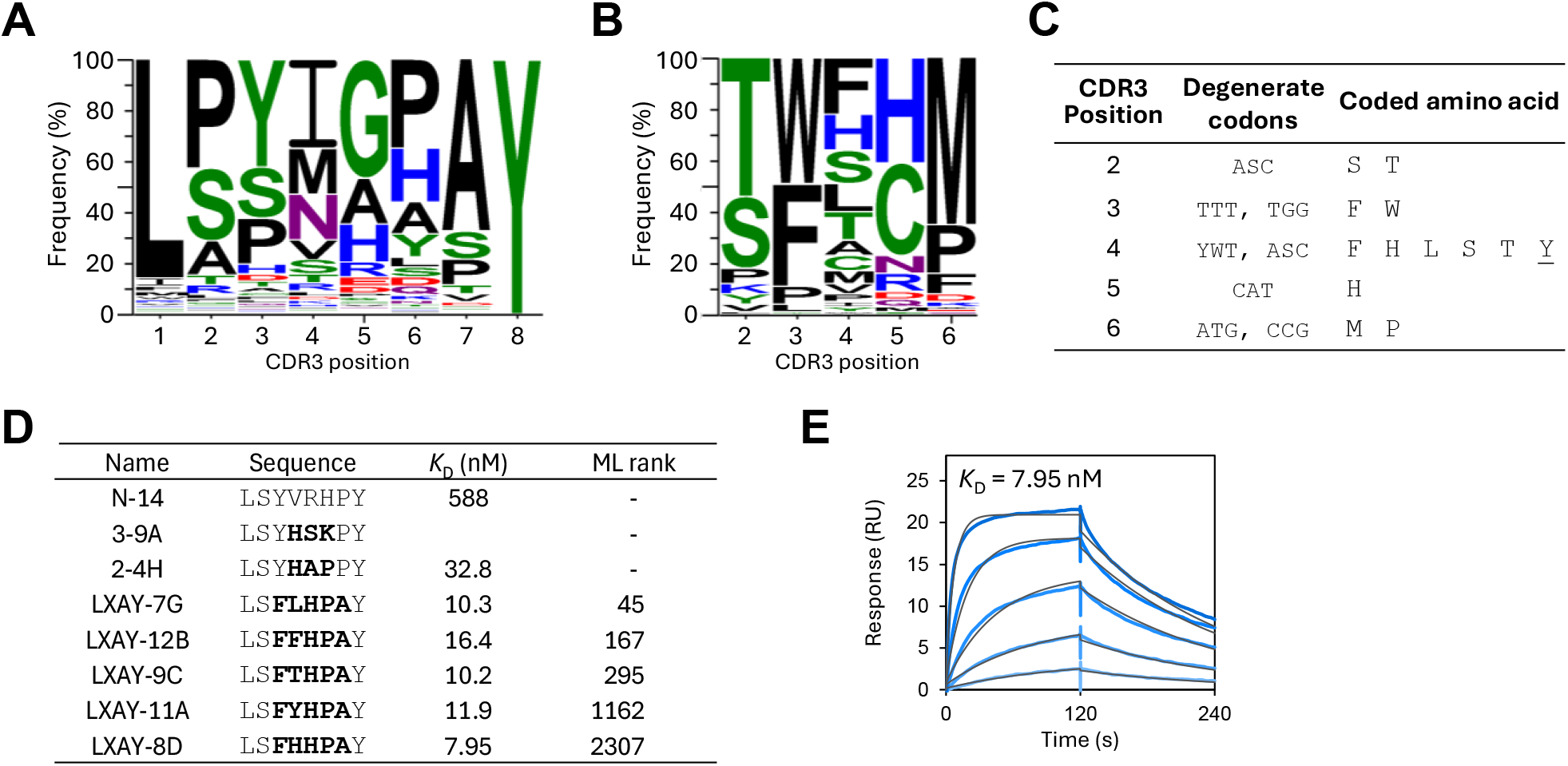
Lead-independent antibody design using machine learning. (A) Amino acid frequencies of variants with scores higher than 1 in the fourth-round PS (positive selection)-eluted phages. (B) Amino acid frequencies at positions 2, 3, 4, 5, and 6 in the ML-predicted top 100 sequences, visualized by using WebLogo. (C) Design of an ML-guided library based on amino acids that appeared with frequencies exceeding 10% in the top 100 sequences predicted by ML. Amino acids not present in these top 100 ML-predicted sequences are underlined. (D) *K*_D_ values and ML-predicted ranks of five high-affinity variants isolated. (E) Kinetic analysis of variant LXAY-8D by surface plasmon resonance. Blue lines represent a three-fold dilution series starting from 133 nM. Black lines indicate the global fit to a 1:1 binding model.

Assuming that these residues were critical for target binding, we finally defined an ML search space restricted to eight-residue CDR3 variants by following the LXXXXXAY motif (X: 20 kinds of amino acids). We extracted sequences matching with the motif from the second-round deep sequencing data, and we obtained 80 sequences for the training data. We trained an ML model to predict the scores of variants in the LXXXXXAY sequence space. Hereafter, this ML is referred to as ML_R2_LX. On the basis of the top 100 predicted sequences (Figure 4B), we designed an ML-guided library incorporating amino acids (excluding cysteine) that appeared with a frequency of 10% or higher at each position (Figure 4C, Table S2). The gene library was transformed into *E. coli*, and 43 randomly selected clones were screened by ELISA. Notably, 20 clones (47%) had binding signals higher than that of N-14 (Figure S3C). Among these, five variants with high binding activity were purified by SEC and subjected to kinetic analysis (Figure 4D). Of these five variants, LXAY-8D (CDR3: LSFHHPAY) showed the strongest binding, with a *K*_D_ value of 7.95 nM (Figure 4E, Figure S5), a 74-fold improvement compared with N-14 and stronger than other affinity-improved variants, including 3-9A and 2-4H. In terms of ML ranking, LXAY-7G was ranked 45th, indicating that the ML model was able to predict high-affinity variants at high ranks.

Collectively, these results indicate that early-round deep sequencing data, which retained a substantial proportion of binding-negative variants, could be effective for identifying high-performance antibodies. They also show that conserved residues in selected libraries could be used to define a lead-independent ML search space, enabling the identification of high-affinity VHHs stronger than those obtained through lead-based maturation.

## Discussion

Here, we aimed to optimize an ML integrated process for functional antibody discovery. Specifically, we investigated the selection of deep sequencing data suitable as training data and validated a lead-independent sequence space design for identifying high-affinity antibodies.

During in vitro selection with phage display, repeated rounds of selection typically lead to the enrichment of binding-positive variants, often resulting in a biased population distribution^5–7^. Here, in contrast, binding-negative variants present in the second round were progressively eliminated, leading to the enrichment of binding-positive variants in the third and fourth rounds (Figure 3F and 3G). This was the ideal result for in vitro selection itself; however, the training data using the second round demonstrated a predictive accuracy superior to that of later rounds for distinguishing between binders and non-binders. This suggested that the presence of binding-negative variants in the second-round dataset might have been a decisive factor in improving model performance. These findings suggest the importance of using training data that include a sufficient representation of non-functional sequences to enhance the success rate of ML-guided antibody discovery. Wu et al.^20^ also suggested the importance of training on a diverse range of sequence data, rather than relying solely on high-performance datasets, for the effective application of ML-based protein-directed evolution. Furthermore, it is a well-established principle in ML that training on imbalanced datasets can substantially degrade predictive performance^21^. Therefore, a dataset dominated only by positive binders may not be optimal for training robust predictive models.

We also designed an ML search space by considering that residues conserved among variants in the fourth round possibly contributed to binding function. By narrowing the search space, ML successfully proposed variants with affinities higher than those obtained through the affinity maturation of N-14. These results demonstrated that optimizing the design of the ML search space allowed us to identify antibodies that had higher binding functions but that might have been missed by lead-dependent approaches. Various other strategies for defining ML search spaces have been reported, such as designing libraries based on high-frequency amino acids at each position^6^, or using deep learning to generate sequences that mimic the properties of selected variants^12^. Further comparative analysis and optimization of sequence space design could further improve both the success rate and the affinity of the identified antibodies.

In recent years, the application of protein language models has shown remarkable results in the field of amino acid sequence embedding^21–23^. These models are trained on hundreds of millions of sequences from databases, enabling the proposal of sequences with optimized structural stability and functional activity. Although we focused here on evaluating how variations in training data influence ML prediction, rather than exploring embedding methods, the use of protein-language-model-based embeddings in our approach could potentially enhance ML accuracy. Furthermore, antibody exploration that combines generative artificial intelligence (AI)-based sequence design with experimental screening is becoming a powerful process for identifying high-performance proteins^24,25^. Although here we used a conventional random library as the initial phage library, integrating AI-designed libraries could improve the probability of obtaining high-affinity variants.

In conclusion, here, we demonstrated that training data selection and ML search space design are critical determinants of successful ML-guided antibody engineering. We showed that early-round deep sequencing data, which retain a substantial proportion of binding-negative variants, can serve as more informative training data than later, more converged datasets. We demonstrated that a lead-independent sequence space designed from residues conserved after in vitro selection enabled the identification of high-affinity variants beyond those obtained through lead-dependent affinity maturation. These findings provide a robust framework for improving the efficiency of ML-guided antibody discovery by using deep sequencing data for both training data selection and search space design, thereby opening up diverse routes for obtaining high-affinity antibodies.

## Methods

### Design of scaffold VHH

From the 268 *Camelus dromedarius* VHH sequences registered in the sdAb-DB^15^, we selected 260 sequences without substantial deletions at either the N- or the C-terminus. These sequences were numbered according to the International Immunogenetics Information System (IMGT) definition^26^, and CDRs were defined by using the AbM system^27,28^. Because of the high variability in CDR3 length, the sequence coverage in CDR3 was relatively low (Figure S1A). The amino acid frequency at each position was calculated (Figure S1B), and a consensus scaffold VHH was designed by adopting the most frequent amino acids at each position (Figure S1C). Cysteine residue at position 38 was replaced with tyrosine, which was the second most frequent amino acid, to prevent unnecessary disulfide bond formation. As a control, a wild-type construct was generated by grafting the CDR3 sequence from PDB (Protein Data Bank) ID 1BZQ (12 residues; GGYELRDRTYGQ) into the consensus scaffold VHH.

### Selection of phages binding to HER3 dimer arm peptide

CDR3-randomized gene fragments were generated by polymerase chain reaction (PCR) using a phagemid vector template containing the wild-type gene and primers incorporating NNB codons. CDR3 lengths were designed to be 8, 12, 16, or 19 residues. The C-terminal residue of CDR3 (position 117) was not mutated, because the frequency of tyrosine at this position is high (65%) (Figure S1B). The resulting mutated VHH gene fragments were cloned into a phagemid vector and transformed into *E. coli* TG-1 cells to generate phage libraries. Phages produced in *E. coli* were precipitated with a polyethylene glycol (PEG)–NaCl solution and subsequently resuspended in phosphate-buffered saline (PBS) (Supporting Methods).

For target preparation, 1 µM HER3 dimer arm peptide, a 21-residue peptide consisting of Biotin-GGGGSLVYNKLTFQLEPNPHT (synthesized by Eurofins Scientific, Tokyo, Japan), was incubated with magnetic beads. For biopanning, the phage library (5 × 10^11^ cfu) was mixed with 10 µL streptavidin-coated magnetic beads (Dynabeads MyOne Streptavidin T1, Thermo Fisher Scientific, Waltham, MA, USA) without antigen for 60 min at room temperature. The magnetic beads and the supernatant were separately collected by magnetic separation (NS in Figure 1C). The supernatant was collected and incubated with HER3-peptide-immobilized magnetic beads for 60 min, and the magnetic beads were subsequently collected (PS in Figure 1C). Next, the magnetic beads from both NS and PS were washed five times with PBS containing 0.05% Tween-20 using a KingFisher purification system (5400000, Thermo Fisher Scientific). The bound phages were then eluted with 100 mM glycine–HCl (pH 2.2) to collect NS-eluted phages from the NS beads and PS-eluted phages from the PS beads. The PS-eluted phages were mixed with 9.8 mL of *E. coli* TG1 culture (OD_600_ = 0.5) and incubated at 37 °C for 60 min for infection. The cells were plated on 2× yeast tryptone (YT) agar medium containing 100 µg/mL ampicillin and 1% (w/v) glucose. Phages were amplified and purified for the next round (amplified phages). Selection was repeated to give a total of four rounds.

### Protein expression

The gene fragments of VHH were amplified from the phagemid vectors by PCR, and the fragments were ligated into the expression vector. Each constructed plasmid, encoding the variant with a FLAG tag and a poly-histidine tag at the C-terminus product, was transformed into *E. coli* BL21(DE3) cells. The cells were incubated overnight at 28 °C on Luria Bertani (LB) agar plates with 100 µg/mL ampicillin. The colonies were transferred to 50 mL LB broth with 100 μg/mL ampicillin in a flask and incubated overnight at 28 °C, and then 5 mL culture was inoculated into 500 mL of 2× YT broth (100 µg/mL ampicillin) in a flask and incubated at 28 °C with shaking. At an OD_600_ of 0.8, IPTG (isopropyl-β-D-thiogalactopyranoside) was added to a final concentration of 1 mM, and the cells were shaken at 28 °C overnight. The cultures were harvested by centrifugation, and the supernatants were collected. Variants were purified from the supernatants by immobilized metal ion affinity chromatography on Ni Sepharose 6 Fast Flow (Cytiva, Marlborough, MA, USA) and by SEC on a HiLoad 26/600 Superdex 75 pg column (Cytiva).

### SPR analysis

The binding affinities of the *E. coli*-expressed VHH variants were evaluated by SPR using a Biacore T200 instrument (Cytiva). Streptavidin was immobilized onto a Series S Sensor Chip CM5 (Cytiva) by amine coupling using an Amine Coupling Kit (Cytiva) for the reference and measurement flow cells. Subsequently, 500 nM biotin was injected into the reference flow cell and 500 nM biotinylated HER3 peptide was injected into the measurement flow cell. Binding assays were conducted by using PBS-T (0.005% Tween 20) as the running buffer at a flow rate of 30 µL/min. The sensor surface was regenerated by using Glycine 2.0 (Cytiva) for 30 s. *K*_D_ values were calculated by global fitting of the 1:1 binding model by using Biacore T200 Evaluation Software (Cytiva).

### Deep sequencing analysis

Following each round of biopanning, ssDNA (single-stranded DNA) was extracted from the initial phages, NS-eluted phages, PS-eluted phages, and amplified phages via phenol– chloroform–isoamyl alcohol extraction. The extracted ssDNAs were amplified by PCR with primers containing adapter sequences, TruSeq DNA CD Indexes (Illumina, San Diego, CA, USA). The PCR products were quantified using a Qbit 1× dsDNA HS Assay Kit (Thermo Fisher Scientific) and pooled in equal amounts. The prepared sample was sequenced on a MiSeq platform (Illumina) using a MiSeq Reagent Kit v3 (Illumina) with 2 × 300-bp paired-end reads. Raw sequencing data were quality filtered, trimmed, and merged. Subsequently, to ensure sequence integrity, only variant sequences of which the sequence except for CDR3 was identical to the wild-type sequence were extracted for further analysis.

### ML prediction

We used an ML framework based on the Bayesian optimization software COMBO, as described previously^6,7,14,29,30^. To construct a training dataset from the deep sequencing data, the performance score of each variant was calculated on the basis of its positive to negative abundance ratio (PS-eluted phages to NS-eluted phages). The performance score of a variant *i* was defined as:

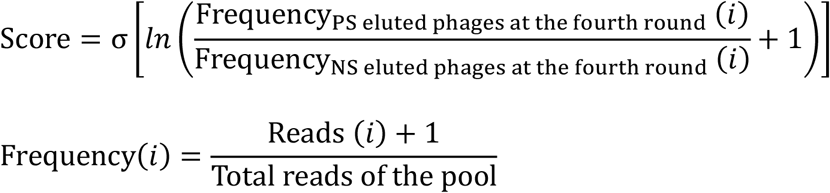

where σ(·) is a sigmoidal function, PS is positive selection, and NS is negative selection. Training datasets were generated by using the deep sequencing data obtained from the second, third, and fourth rounds. Specifically, we extracted variants with a read count of 4 or more in either the PS or the NS phages. This yielded 6771 sequences for the second round, 3146 sequences for the third round, and 871 sequences for the fourth round (Figure S2). The feature vector of a VHH variant was defined by concatenating the precomputed feature vectors of amino acids at the mutated sites (CDRs). To select the optimal feature vector of each amino acid, we tested a variety of amino acid descriptors, as previously described^6^, and we found that the ProtFP descriptor^31^ provided the best accuracy for the third-round data. The ProtFP descriptor was used to construct ML models with other training data.

The trained models were used to rank all variants within these predicted sequence spaces according to their probability-of-improvement scores. Following the ML process, mutation rates for the top-ranked sequences were calculated by determining the proportion of variants with a mutation at each residue position. The ML-predicted top 100 sequences were visualized using WebLogo^32^.

### Protein expression in 96-well plates

The gene fragments of the ML-guided VHH library were generated from plasmids containing functional VHH sequences by overlap extension PCR using primers with degenerate codons. They were then ligated into the expression vector. *E. coli* BL21(DE3) cells were transformed with the resultant vectors and incubated on LB agar plates (100 µg/mL ampicillin) overnight at 28 °C. The colonies were randomly transferred to 1 mL LB broth (100 μg/mL ampicillin) in a deep-well plate (Axygen, Union City, CA, USA) and incubated overnight at 28 °C. Then, 10 µL of the culture was inoculated into 990 µL of 2× YT broth (100 µg/mL ampicillin) in another deep-well plate and incubated at 28 °C with shaking. At a culture OD_600_ of 0.8, IPTG was added to a final concentration of 1 mM, and the cells were shaken at 28 °C overnight. The cultures were harvested by centrifugation, and the supernatants were used for ELISA.

### ELISA

A 96-well plate was coated with 50 μL of 4 μg/mL NeutrAvidin (Thermo Fisher Scientific) in PBS for 60 min at room temperature, and 150 µL of 0.5% (w/v) bovine serum albumin in PBS was added for 30 min for blocking. After each well had been washed with PBS, 50 µL of 500 nM biotinylated HER3 dimer-arm peptide was added and the plates were incubated for 30 min. PBS was used in the reference wells. Each well was washed with PBS, followed by incubation with the supernatant of the VHH variants for 30 min. Each well was washed three times with PBS containing 0.05% Tween-20, and 50 µL of 10,000-fold diluted horseradish peroxidase-conjugated anti-FLAG tag monoclonal antibody (A8592, Sigma Aldrich, St. Louis, MO, USA) was added to each well and incubated for 40 min. Each well was washed as above, and 50 μL of 3,3′,5,5′-tetramethylbenzidine solution (1-step Ultra TMB-ELISA Substrate Solution, Thermo Fisher Scientific) was added. The plate was incubated for 10 min at room temperature. After the incubation, 50 μL of 2 M H_2_SO_4_ was added to each well and the absorbance at 450 nm was measured with a Synergy H4 Hybrid Multimode Microplate Reader (BioTek, Winooski, VT, USA)

## Acknowledgments

The authors thank Ms. Hiromi Ito, Ms. Miho Hosoya, and Ms. Yuri Ishigaki for experimental support. This work was supported partly by the projects “Development of the Key Technologies for the Next-Generation Artificial Intelligence/Robots” (M.U., 20000157-0) and “Development of Quantum-Classical Hybrid Use-Case Technologies in Cyber-Physical Space” of the New Energy and Industrial Technology Development Organization (M.U., 23201802-0); by a Scientific Research Grant from the Japan Society for the Promotion of Science research fellowship (M.U., JP20H00315, JP24K01260); by a “Fundamental drug discovery technology development project to realize next-generation treatments and diagnosis” of the Japan Agency for Medical Research and Development (M.U., 23ae0121003h0003); by the Japan Society for the Promotion of Science (S.K., 22KJ0218); and by Support for Pioneering Research Initiated by the Next Generation (SPRING) from the Japan Science and Technology Agency (S.K., JPMJFS2102; T.I., JPMJSP2114).

## Author information

T.I., S.K., and M.U. conceived the research strategy; T.I., S.K., and H.N. developed the methodology; T.I., S.K., Y.K., and Y.S. contributed to the design of the ML search spaces; S.K. performed experiments and visualized the experimental data; and T.I. and M.U. wrote the original draft of the manuscript.

## Data availability

The datasets used in this study are available from the corresponding author on request.

