## Supplementary Information for "Exploring diverse routes to high-affinity-antibody variable domains through deep-sequencing-informed machine learning"

[1] Department of Biomolecular Engineering, Graduate School of Engineering, Tohoku University, Sendai, Japan; [2] Department of Data Science, School of Frontier Engineering, Kitasato University, Kanagawa, Japan; [3] Artificial Intelligence Research Center, National Institute of Advanced Industrial Science and Technology (AIST), Tokyo, Japan; [4] Department of Computational Biology and Medical Sciences, Graduate School of Frontier Sciences, The University of Tokyo, Chiba, Japan; [5] Center for Advanced Intelligence Project, RIKEN, Tokyo, Japan

### Both authors contributed equally to this work.

#### Supporting Methods

##### *Phage preparation*

The mutated VHH gene fragments were cloned into a phagemid vector and transformed into *Escherichia coli* TG-1 (Lucigen) to generate phage libraries. The transformants were plated on 2× YT agar medium containing 100 µg/mL ampicillin and 1% (w/v) glucose, followed by overnight incubation at 37 °C. Colonies were scraped off using a spreader and inoculated into 2× YT medium containing 100 µg/mL ampicillin and 1% (w/v) glucose to an initial $OD_{600}$ of 0.1. The culture was incubated at 37 °C with shaking at 200 rpm until the $OD_{600}$ reached 0.5. Subsequently, the culture was infected with the helper phage M13KO7 ($2 \times 10^{12}$ pfu/mL; NEB Japan Inc., Tokyo, Japan) and incubated without shaking at 37 °C for 60 min. Cells were then collected by centrifugation and incubated overnight at 28 °C and 100 rpm in 2× YT medium supplemented with 100 µg/mL ampicillin, 50 µg/mL kanamycin, and 1 mM IPTG. After incubation, the culture supernatant was collected by centrifugation, mixed with a 25% PEG 6000–2.5 M NaCl solution, and incubated on ice for 1 h. Phage particles were precipitated by centrifugation and the resulting pellet was resuspended in 1 mL PBS. The phage library was further purified by repeating the PEG–NaCl precipitation and resuspension steps.

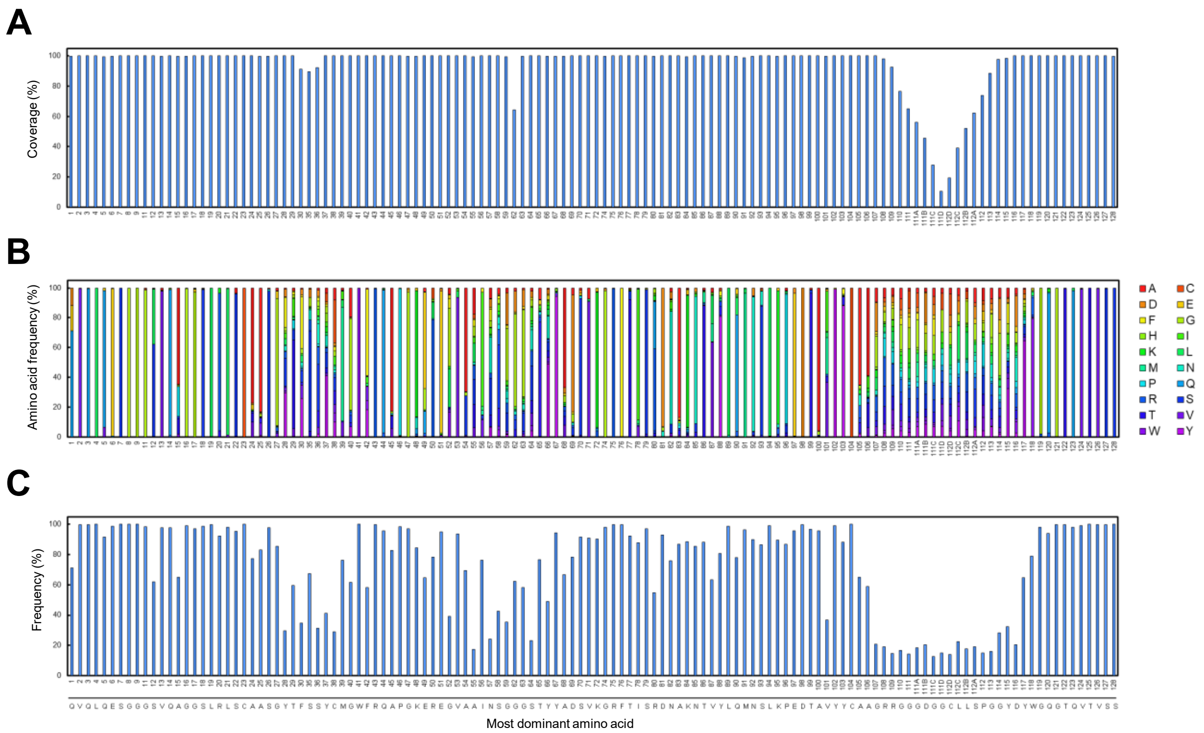


**Figure S1. Sequence profiling and amino acid conservation of VHHs.**

(A) Sequence coverage at each position according to the IMGT numbering scheme. (B) Amino acid frequency distribution at each position. (C) Frequency of the most dominant amino acid at each position.

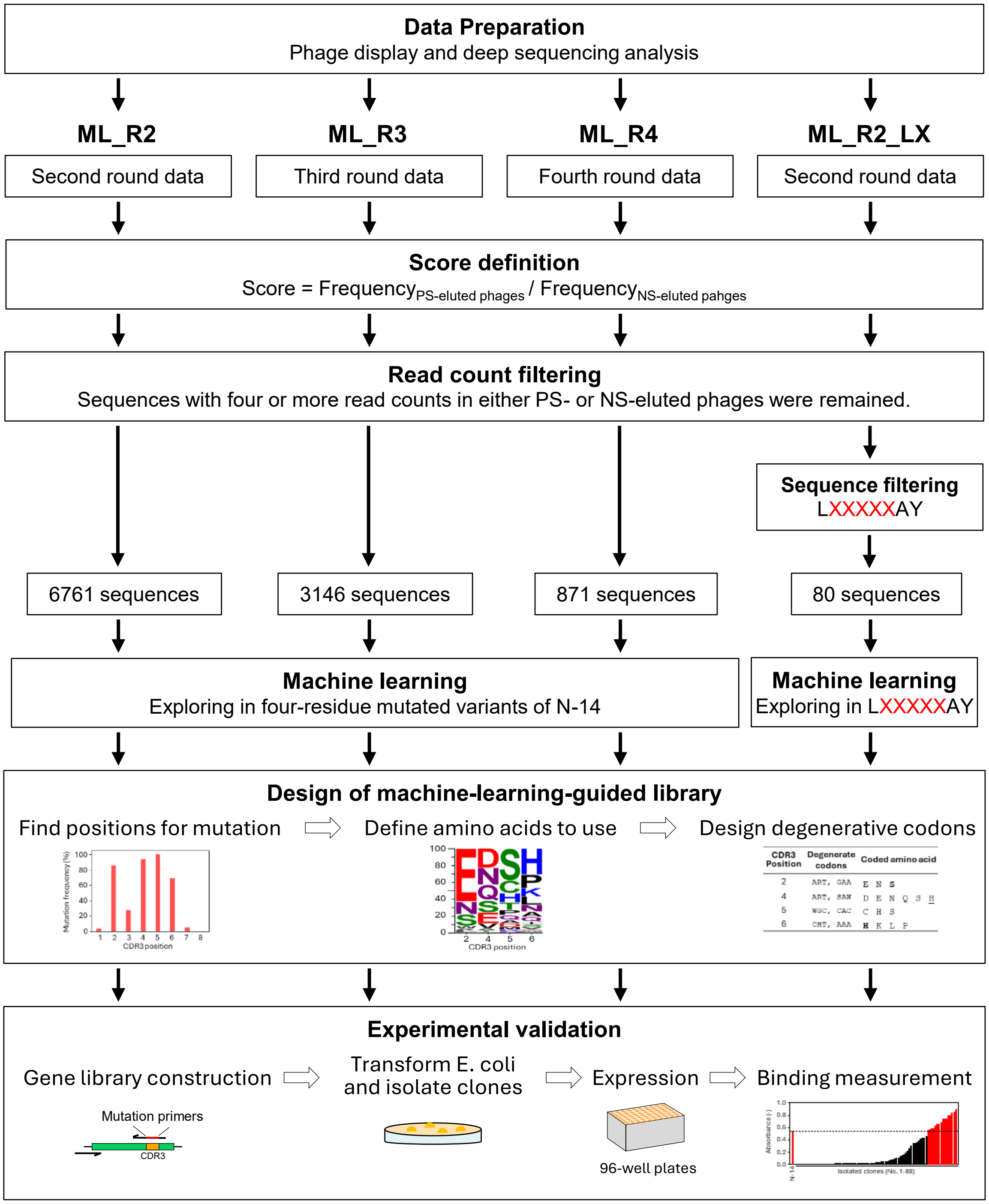


**Figure S2. Workflow of machine-learning-guided VHH identification**

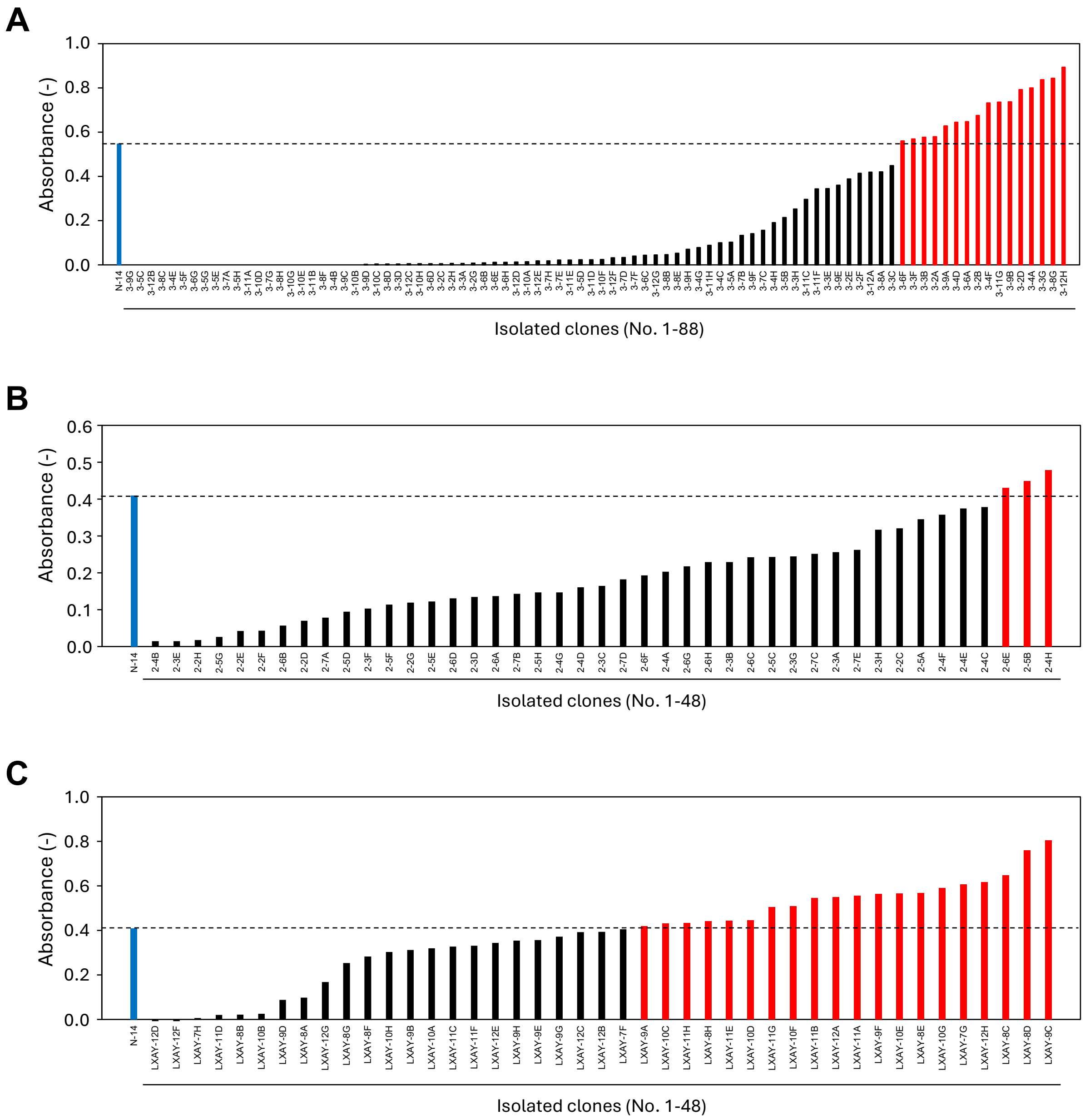


**Figure S3. Evaluation of binding activity for machine-learning-guided libraries.** Monoclonal ELISA for the ML_R3-guided library (A), the ML_R2-guided library (B), and the ML_R2_LX-guided library (C). In all figures, the lead molecule N-14 is indicated in blue, whereas clones with binding signals higher than that of N-14 are in red.

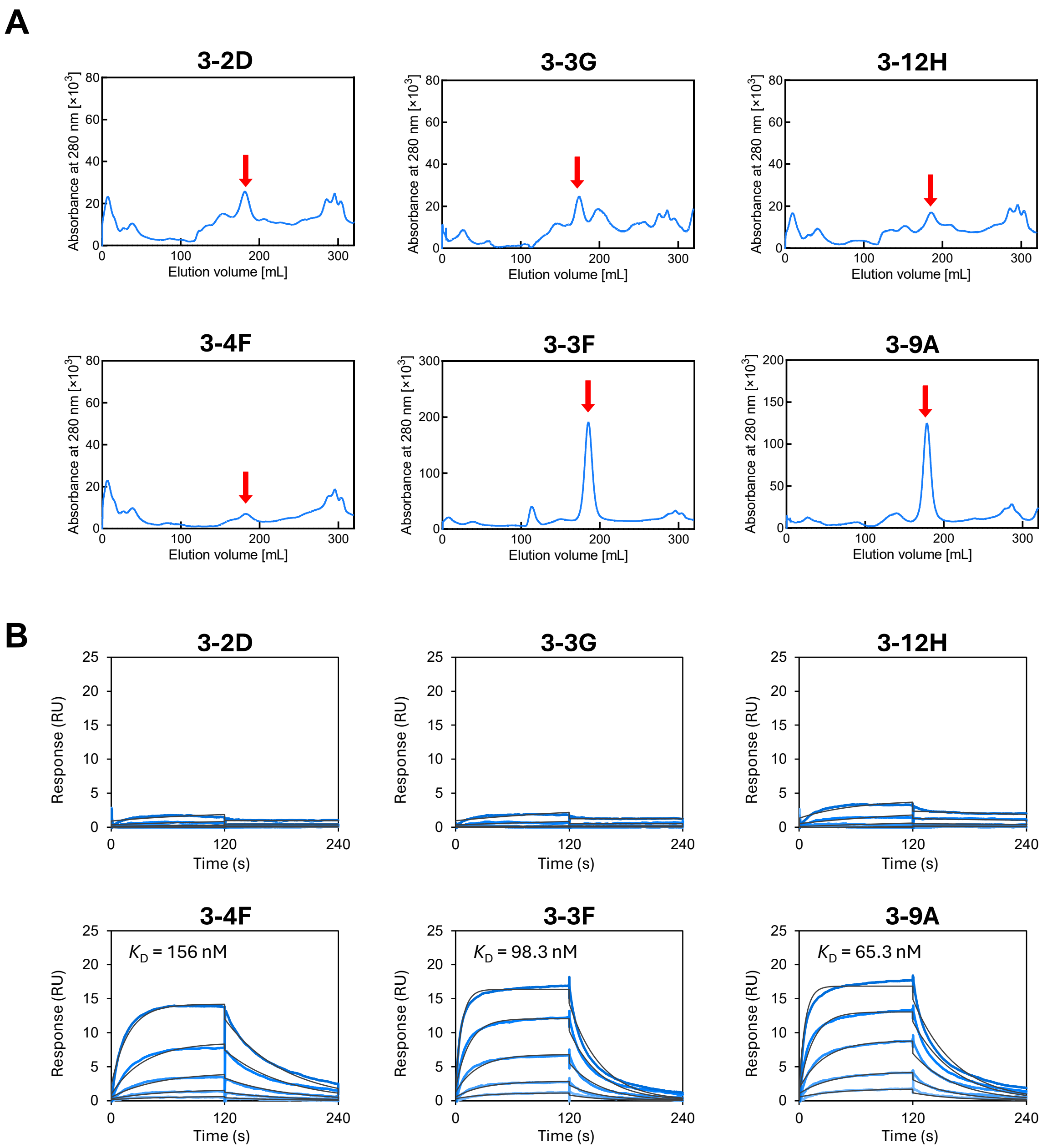


**Figure S4. Evaluation of variants identified from the ML_R3-guided library.**
(A) Size-exclusion chromatograph on a HiLoad 26/600 Superdex 75 pg (Cytiva). The absorbance of the eluent was monitored at a wavelength of 280 nm. Red arrows indicate collected fractions used for further analysis. (B) Kinetic analysis with surface plasmon resonance. Three-fold dilution series were used for the analytes, starting from 400 nM (blue lines). $K_D$ values were determined by global fitting using a 1:1 binding model (black lines).

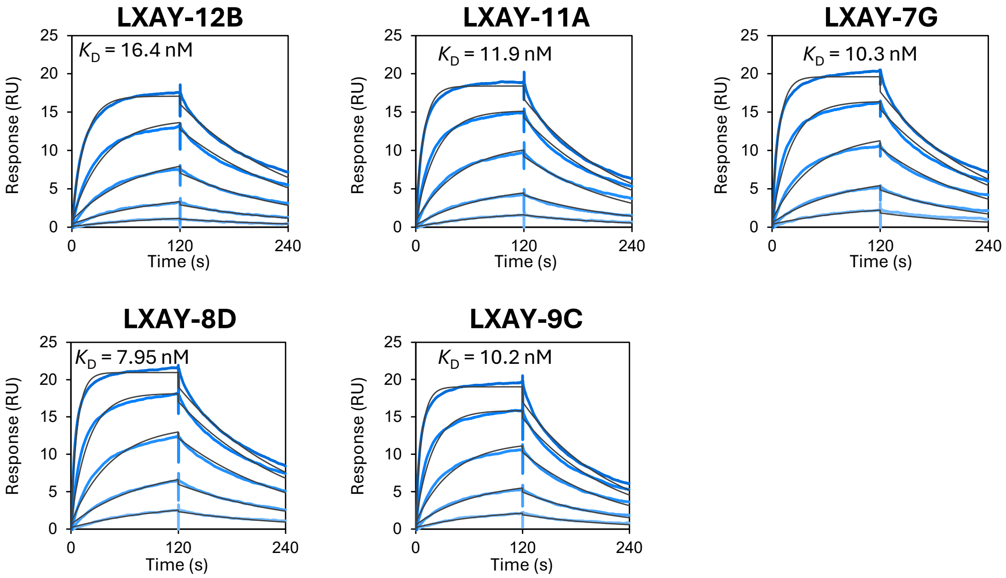


**Figure S5. Evaluation of variants identified from the ML_R3-guided library.** Kinetic analysis with surface plasmon resonance. Three-fold dilution series were used for the analytes, starting from 133 nM (blue lines). $K_D$ values were determined by global fitting using a 1:1 binding model (black lines).

#### Table S1. Machine-learning-predicted top 100 sequences.

| | Predicted sequences | | |
|---|---|---|---|
| RANK | ML_R2 | ML_R3 | ML_R4 |
| 1 | LSYHMPPY | LNYQCLPY | LSYMAPAY |
| 2 | LSYNMPPY | LEFSSHPY | LSYDEMAY |
| 3 | LAYERCPY | LEYDSHPY | LSYDAMAY |
| 4 | LSYHFPPY | LEFNSHPY | ESYDEMPY |
| 5 | LVYNMPPY | LNYNCLPY | LSYDAPAY |
| 6 | LVYHMPPY | LEYSSHPY | LSYSEPPY |
| 7 | LAYEQCPY | LNYECLPY | LSYDEMPY |
| 8 | LSYSAKPY | LEYNSNPY | LSYSDPPY |
| 9 | LTYHMPPY | LEFDSHPY | LSYAEPPY |
| 10 | LSWNAPPY | LEYNSHPY | LTYSDPPY |
| 11 | LSYNAPPY | LEYDSQPY | ESYDEHPY |
| 12 | LQYHFPPY | LEYDSHAY | LTYSEPPY |
| 13 | LVYHFPPY | LEYQCLPY | LSYMAPPY |
| 14 | LSWNQSPY | LEFTSHPY | LTYDEMPY |
| 15 | LSWNASPY | LEYSSQPY | LSYASPPY |
| 16 | LSHEEHLY | LSYEHPPY | LSYVSPPY |
| 17 | LSWHMPPY | LSYEMPPY | LSYISPPY |
| 18 | LSWHAPPY | LEYNSQPY | LDYDDPPY |
| 19 | LSYNFPPY | LEFSTHPY | LSYSAPPY |
| 20 | LSFNMPPY | LEYDSLPY | ENYVEHPY |
| 21 | LVYEMCPY | LEFNTHPY | LSYAAPAY |
| 22 | LSYQAKPY | LEYSSNPY | LDYSDPPY |
| 23 | LTYNMPPY | LNYQCVPY | LSYVDPPY |
| 24 | LSWNATPY | LNYQCIPY | LSDSDPPY |
| 25 | LSWNMPPY | LEYDTHPY | ESYSEPPY |
| 26 | LTYHFPPY | PWNVDHPY | LSYDEPPY |
| 27 | LSWNSSPY | LEWSSHPY | ENYIEHPY |
| 28 | LQYHMPPY | LEYQSKPY | LSDSSPPY |
| 29 | LSWNQTPY | LNYEMPPY | LTYDSPPY |
| 30 | LSHEEHFY | LEFQSHPY | LTYDEPPY |
| 31 | LSYHAPPY | LEYNSSPY | LSFSDPPY |
| 32 | LSYNQPPY | LEYDSNPY | LSYVEPPY |
| 33 | LVYNPSPY | LEFNDHPY | LSYSSPPY |
| 34 | LNYNFPPY | LSIQHPPY | LTYSSPPY |
| 35 | LTYNAPPY | LSYEHPAY | LTYMAPPY |
| 36 | LLYERCPY | LEYSSSPY | ESYSDPPY |
| 37 | LGYNFHFY | LLYNSNPY | LTYISPPY |
| 38 | LNYNMPPY | LNYECPPY | LTYVSPPY |
| 39 | LAWERCPY | LEYDQHPY | ETYDEHPY |
| 40 | LSFNAPPY | LEYSSKPY | LTYAEPPY |
| 41 | LQYNMPPY | LEYQPKPY | LTYVDPPY |
| 42 | LSYHLPPY | LSYQHPPY | LSDSAPPY |
| 43 | LLYEQCPY | LEYSSAPY | LTYDAHAY |
| 44 | LLYEMTPY | LEFDTHPY | LTYSAPPY |
| 45 | LGYNFHAY | ISYEHPPY | LSYTDPPY |
| 46 | LTYHAPPY | LEYDSSPY | LDYDSPPY |
| 47 | LLYEWCPY | LSIEHPPY | LDYSEPPY |
| 48 | LNYCCPPY | LAEVKNPY | LSYIDPPY |
| 49 | LVYHCPPY | LNYQCAPY | LTYTDPPY |
| 50 | LVYHSPPY | LSYQHPAY | LSYDAPPY |
| 51 | INAVGHPY | LNYNCVPY | LDYDEPPY |
| 52 | LSYNLHPY | LEYDSAPY | LTYDAPPY |
| 53 | LSTEEMPY | LEYQSQPY | LSYDSPPY |
| 54 | LVYNFPPY | LWDVNKPY | LSYVNPPY |
| 55 | LSWNQPPY | LNYNCIPY | LDYDAPPY |
| 56 | LLYEQTPY | LAEVKDPY | LSYAAPPY |
| 57 | LNYHFPPY | LEYDSKPY | LSYSPPPY |
| 58 | LSWNFTPY | LSDASKPY | LSISEPPY |
| 59 | LSYNASPY | LEFDQHPY | EDYVDPPY |
| 60 | LTYHLPPY | LEYDSMPY | LTYDEHAY |
| 61 | LLYEFTPY | LEFNQHPY | LSIVDPPY |
| 62 | LSIGWEPY | LNYQHPPY | LSYDEHAY |
| 63 | LGSVWHLY | LEYDPHPY | LSHDEMPY |
| 64 | LVYQPSPY | LEYDAHPY | LSYDASAY |
| 65 | LEFERTPY | LNYEHPPY | LSYAPPPY |
| 66 | LSHEQYPY | PSNVMPPY | LSYAEPAY |
| 67 | LVYEFCPY | LEYNTHPY | LSYTEPPY |
| 68 | LNAVGHLY | LEYESKPY | LTYIDPPY |
| 69 | LSWNFSPY | LSVQHPPY | LTYASPPY |
| 70 | LSHENCPY | LNYECIPY | LSIVNPPY |
| 71 | LSWNNGPY | LEWNSHPY | LTYDDPPY |
| 72 | LSVNFPPY | LEYSSLPY | LSDSMPPY |
| 73 | INSVGHPY | LEYNQNPY | LSISDPPY |
| 74 | LSYHCPPY | PFNVDHPY | LSYINPPY |
| 75 | VSYNMPPY | LEYSPKPY | ESYVDPPY |
| 76 | LSYHQPPY | LEYDSRPY | ESYAEPPY |
| 77 | LLYEMSPY | LEYTSHPY | LSTSDPPY |
| 78 | LSYSQKPY | LEYQSHPY | ESYDEPPY |
| 79 | LSWNSTPY | LEFSQHPY | LSIIDPPY |
| 80 | LSQEEMPY | LNYNCAPY | LSYDAHAY |
| 81 | LVYHAPPY | LEYQSLPY | LSYIEPPY |
| 82 | LDYNLPPY | LEFSAHPY | LSIINPPY |
| 83 | LSHEEHIY | LEYDPAPY | LSYADPPY |
| 84 | LLYEFCPY | LEYNANPY | LNYVEHPY |
| 85 | LNYNQPPY | LEYNTNPY | LSDDSPPY |
| 86 | LVYEQCPY | LWYCNGPY | ETYVDPPY |
| 87 | LSFNASPY | LSYEHPTY | LSNDEMPY |
| 88 | DSGVGHIY | LEYNQHPY | LTYSPPPY |
| 89 | LLYEWFPY | LNYECVPY | LSYVHPPY |
| 90 | LSYNFSPY | LEYDSHCY | LSIVEPPY |
| 91 | LQYHQPPY | LSDATKPY | LNYIEHPY |
| 92 | LNYNAPPY | LEFQDHPY | LSYDDPPY |
| 93 | LSHEEHVY | LLYQPAPY | LSYMEPPY |
| 94 | LSYHFSPY | LEYDTLPY | LSYEAPAY |
| 95 | LQYHAPPY | LEYNSKPY | LSYSAPAY |
| 96 | ASYHCPPY | LEYSCLPY | LDYTDPPY |
| 97 | LSHECAPY | LSFQCLPY | LSYDEHPY |
| 98 | LNLNAHPY | LEYDAPPY | LSNDEPPY |
| 99 | LSWHCSPY | LEYDTQPY | LSYANPPY |
| 100 | LVYEMSPY | LEYDPQPY | LSVSDPPY |

**Table S2. Amino acid frequencies of machine-learning-predicted sequences.** Amino acid frequencies (%) in the top 100 sequences predicted by models ML_R2, ML_R3, and ML_R2_LX. Amino acids with frequencies of 10% or higher are indicated in red text, and those consistent with N-14 are surrounded by blue squares.

**ML_R2**

| | 1 | 2 | 3 | 4 | 5 | 6 | 7 | 8 |
|---|---|---|---|---|---|---|---|---|
| A | 1 | 3 | 2 | 0 | 17 | 1 | 1 | 0 |
| C | 0 | 0 | 0 | 1 | 6 | 11 | 0 | 0 |
| D | 1 | 1 | 0 | 0 | 0 | 0 | 0 | 0 |
| E | 0 | 1 | 0 | 26 | 6 | 1 | 0 | 0 |
| F | 0 | 0 | 4 | 0 | 18 | 1 | 2 | 0 |
| G | 0 | 3 | 1 | 1 | 4 | 1 | 0 | 0 |
| H | 0 | 0 | 7 | 25 | 0 | 13 | 0 | 0 |
| I | 2 | 0 | 1 | 0 | 0 | 0 | 2 | 0 |
| K | 0 | 0 | 0 | 0 | 0 | 3 | 0 | 0 |
| L | 95 | 9 | 1 | 0 | 4 | 0 | 3 | 0 |
| M | 0 | 0 | 0 | 0 | 17 | 2 | 0 | 0 |
| N | 0 | 10 | 0 | 38 | 2 | 0 | 0 | 0 |
| P | 0 | 0 | 0 | 0 | 2 | 45 | 91 | 0 |
| Q | 0 | 5 | 1 | 2 | 13 | 0 | 0 | 0 |
| R | 0 | 0 | 0 | 0 | 4 | 0 | 0 | 0 |
| S | 0 | 49 | 2 | 2 | 3 | 13 | 0 | 0 |
| T | 0 | 6 | 1 | 0 | 0 | 8 | 0 | 0 |
| V | 1 | 13 | 1 | 5 | 0 | 0 | 1 | 0 |
| W | 0 | 0 | 16 | 0 | 4 | 0 | 0 | 0 |
| Y | 0 | 0 | 63 | 0 | 0 | 1 | 0 | 100 |

**ML_R3**

| | 1 | 2 | 3 | 4 | 5 | 6 | 7 | 8 |
|---|---|---|---|---|---|---|---|---|
| A | 0 | 2 | 0 | 2 | 4 | 6 | 3 | 0 |
| C | 0 | 0 | 0 | 1 | 15 | 0 | 1 | 0 |
| D | 0 | 0 | 3 | 23 | 4 | 1 | 0 | 0 |
| E | 0 | 63 | 2 | 13 | 0 | 0 | 0 | 0 |
| F | 0 | 1 | 15 | 0 | 0 | 0 | 0 | 0 |
| G | 0 | 0 | 0 | 0 | 0 | 1 | 0 | 0 |
| H | 0 | 0 | 0 | 0 | 11 | 31 | 0 | 0 |
| I | 1 | 0 | 2 | 0 | 0 | 3 | 0 | 0 |
| K | 0 | 0 | 0 | 0 | 2 | 10 | 0 | 0 |
| L | 96 | 2 | 0 | 0 | 0 | 10 | 0 | 0 |
| M | 0 | 0 | 0 | 0 | 3 | 1 | 0 | 0 |
| N | 0 | 15 | 3 | 20 | 2 | 8 | 0 | 0 |
| P | 3 | 0 | 0 | 0 | 6 | 16 | 95 | 0 |
| Q | 0 | 0 | 0 | 19 | 6 | 6 | 0 | 0 |
| R | 0 | 0 | 0 | 0 | 0 | 1 | 0 | 0 |
| S | 0 | 14 | 0 | 14 | 38 | 3 | 0 | 0 |
| T | 0 | 0 | 0 | 2 | 9 | 0 | 1 | 0 |
| V | 0 | 0 | 1 | 6 | 0 | 3 | 0 | 0 |
| W | 0 | 3 | 2 | 0 | 0 | 0 | 0 | 0 |
| Y | 0 | 0 | 72 | 0 | 0 | 0 | 0 | 100 |

**ML_R2_LX**

| | 1 | 2 | 3 | 4 | 5 | 6 | 7 | 8 |
|---|---|---|---|---|---|---|---|---|
| A | 0 | 0 | 0 | 6 | 0 | 1 | 100 | 0 |
| C | 0 | 0 | 0 | 6 | 36 | 0 | 0 | 0 |
| D | 0 | 0 | 0 | 0 | 3 | 3 | 0 | 0 |
| E | 0 | 0 | 0 | 0 | 0 | 1 | 0 | 0 |
| F | 0 | 0 | 40 | 22 | 0 | 8 | 0 | 0 |
| G | 0 | 0 | 0 | 0 | 0 | 0 | 0 | 0 |
| H | 0 | 0 | 0 | 14 | 41 | 0 | 0 | 0 |
| I | 0 | 0 | 0 | 2 | 0 | 0 | 0 | 0 |
| K | 0 | 4 | 0 | 0 | 0 | 2 | 0 | 0 |
| L | 100 | 1 | 3 | 11 | 0 | 0 | 0 | 0 |
| M | 0 | 0 | 0 | 5 | 2 | 65 | 0 | 0 |
| N | 0 | 0 | 0 | 0 | 7 | 0 | 0 | 0 |
| P | 0 | 6 | 7 | 3 | 0 | 19 | 0 | 0 |
| Q | 0 | 0 | 0 | 0 | 3 | 1 | 0 | 0 |
| R | 0 | 0 | 0 | 0 | 7 | 0 | 0 | 0 |
| S | 0 | 28 | 0 | 13 | 1 | 0 | 0 | 0 |
| T | 0 | 54 | 0 | 11 | 0 | 0 | 0 | 0 |
| V | 0 | 3 | 0 | 4 | 0 | 0 | 0 | 0 |
| W | 0 | 0 | 49 | 1 | 0 | 0 | 0 | 0 |
| Y | 0 | 4 | 1 | 2 | 0 | 0 | 0 | 100 |

**Table S3. Amino acid sequences and machine-learning ranks of high-binding clones isolated from the ML_R3-guided library.**

| Name | Position | | | | ELISA rank | ML_R3 predicted rank |
|---|---|---|---|---|---|---|
| | 2 | 4 | 5 | 6 | | |
| N-14 | S | V | R | H | 17 | 239817 |
| 3-12H | S | N | C | L | 1 | 3007 |
| 3-8G | S | D | C | L | 2 | 1826 |
| 3-3G | S | D | C | L | 3 | 1826 |
| 3-4A | S | H | H | H | 4 | 919218 |
| 3-2D | S | E | C | L | 5 | 338 |
| 3-9B | S | N | C | L | 6 | 3007 |
| 3-11G | S | H | H | H | 7 | 919218 |
| 3-4F | S | S | H | P | 8 | 4561 |
| 3-2B | S | E | C | L | 9 | 338 |
| 3-6A | S | H | S | K | 10 | 7860 |
| 3-4D | S | H | S | K | 11 | 7860 |
| 3-9A | S | H | S | K | 12 | 7860 |
| 3-2A | S | S | C | L | 13 | 10951 |
| 3-3B | S | N | H | H | 14 | 166867 |
| 3-3F | S | S | S | K | 15 | 6682 |
| 3-6F | S | S | S | L | 16 | 58079 |

**Table S4. Amino acid sequences and machine-learning ranks of high-binding clones isolated from the ML_R2-guided library.**

| Name | Position | | | ELISA rank | ML_R2 predicted rank |
|---|---|---|---|---|---|
| | 4 | 5 | 6 | | |
| N-14 | V | R | H | 4 | 393001 |
| 2-4H | H | A | P | 1 | 31 |
| 2-5B | H | Q | P | 2 | 76 |
| 2-6E | H | Q | H | 3 | 33032 |
